# A PLP-Dependent Decarboxylative Mannich Reaction Initiates Construction of the Nonpeptidic Scaffold of Kaitocephalin

**DOI:** 10.64898/2026.06.22.733665

**Authors:** Tomohiro Noguchi, Yukari Maeno, Kazuo Shin-ya, Tomohisa Kuzuyama

## Abstract

Kaitocephalin (KCP) is a fungal neuroactive natural product bearing a peptide-like yet nonpeptidic amino acid-derived scaffold in which amino acid-like units are connected by C–C bonds rather than peptide bonds. The enzymatic construction of this unusual scaffold has remained unresolved. Here, we identify KpbH as a PLP-dependent enzyme that couples pyrroline-5-carboxylate, generated from L-ornithine, with L-aspartate to form (2*S*,5*R*)-5-((*S*)-2-amino-2-carboxyethyl)pyrrolidine-2-carboxylic acid (ACPCA), which corresponds to the nonpeptidic Ala–Pro substructure of KCP. D_2_O-labeling experiments showed enzyme-controlled, solvent-derived deuterium incorporation at C7 of ACPCA, supporting a decarboxylative Mannich-type mechanism. Feeding of a deuterium-enriched ACPCA-containing reaction mixture to the KCP-producing fungus *Eupenicillium shearii* resulted in deuterium incorporation into KCP, linking ACPCA to KCP biosynthesis. These results identify KpbH as the first native PLP-dependent enzyme that catalyzes an L-aspartate-dependent decarboxylative Mannich-type C–C bond-forming reaction and reveal a biosynthetic strategy for constructing a noncanonical amino acid-like C–C bond scaffold.

**GRAPHICAL ABSTRACT:** 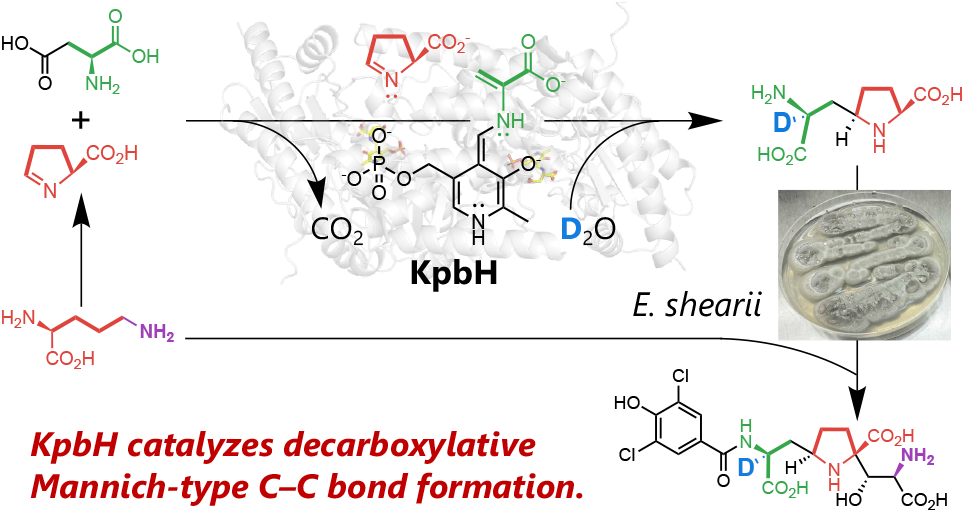

---

Kaitocephalin (KCP), a neuroactive fungal natural product isolated from *Eupenicillium shearii* PF1191, comprises L-alanine-, L-proline-, and D-serine-like units together with an amide-linked 3,5-dichloro-4-hydroxybenzoyl moiety (Figure 1).^1,2^ Although the three amino acid-like units make KCP appear peptide-like at first glance, they are connected through two C–C bonds rather than peptide bonds, resulting in an entirely nonpeptidic amino acid-derived scaffold. This unusual connectivity cannot be readily explained by typical peptide, polyketide, or terpenoid biosynthetic logic, which had long hampered identification of the corresponding biosynthetic gene cluster.^3–6^

**Figure 1.**
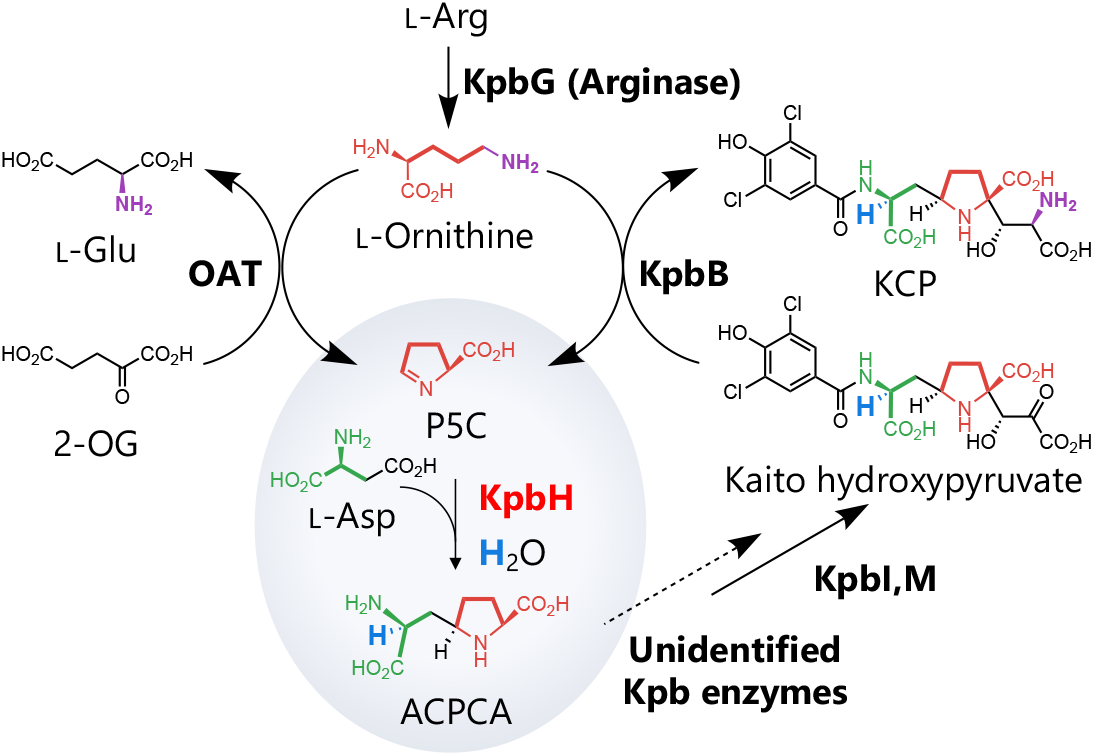
Proposed pathway for construction of the nonpeptidic Ala–Pro-like scaffold of KCP. OAT generates P5C from L-ornithine, and KpbH couples P5C with L-aspartate through a PLP-dependent decarboxylative Mannich-type C–C bond-forming reaction to form ACPCA with incorporation of a solvent-derived hydrogen/deuterium atom. A dashed arrow indicates proposed downstream transformations from ACPCA for which the responsible Kpb enzymes remain unidentified, preceding the KpbI/KpbM-catalyzed steps and KpbB-catalyzed transamination that yield KCP. In the KpbB reaction, L-ornithine serves as the amino donor and is converted to P5C. Colors indicate proposed origins: red, P5C; green, L-aspartate; purple, L-ornithine-derived amino group; blue, solvent-derived hydrogen/deuterium.

We recently identified the KCP biosynthetic gene cluster, designated the *kpb* cluster (Figure S1), and characterized several late-stage biosynthetic intermediates, which enabled biochemical assignment of the α-ketoglutarate-dependent dioxygenase KpbI, the halogenase KpbM, and the aminotransferase KpbB as enzymes responsible for late-stage transformations, including formation of the D-serine moiety (Figure 1).^7^ In contrast to these late-stage transformations, our intermediate analyses suggested that formation of the two nonpeptidic C–C bonds occurs at an early stage of KCP biosynthesis. However, the responsible enzymes and mechanisms remained unknown.

The previous observation that KCP production is enhanced by the addition of adenine and L-arginine, together with [^13^C_6_]-L-arginine feeding experiments showing incorporation of labeled carbons into the proline-like unit of KCP, suggested that an arginine-derived L-ornithine/pyrroline-5-carboxylate (P5C) pathway supplies this part of the scaffold.^7^ However, the enzymatic step that couples this P5C-derived unit to the apparent alanine-like portion of KCP, thereby forming the nonpeptidic Ala–Pro substructure, had not been identified.

Here, we show that KpbH, a pyridoxal 5′-phosphate (PLP)-dependent enzyme encoded in the *kpb* cluster, catalyzes decarboxylative Mannich-type C–C bond formation between P5C and L-aspartate to generate the nonpeptidic Ala–Pro substructure of KCP. This finding identifies a native PLP-dependent enzyme that catalyzes an L-aspartate-dependent decarboxylative Mannich-type C–C bond-forming reaction and provides a biochemical basis for construction of the unusual nonpeptidic scaffold of KCP.

PLP-dependent enzymes are central catalysts in amino acid metabolism and have increasingly been recognized as versatile enzymes for C–C bond formation in natural product biosynthesis.^8–10^ Among these transformations, decarboxylative reactions using L-aspartate-derived nucleophiles are particularly relevant to the formation of the nonpeptidic Ala–Pro substructure of KCP, because they convert a simple amino acid into a reactive carbon nucleophile. UstD provides a representative example, catalyzing PLP-dependent decarboxylative aldol reactions through generation of an L-aspartate-derived nucleophilic intermediate.^11–14^ Recently, this UstD-type catalytic logic was extended to engineered UstD-catalyzed Mannich reactions using imine electrophiles, enabling the synthesis of γ-chiral amino-substituted α-amino acids (Figure S2).^15– 18^ However, despite these advances in engineered biocatalysis, native PLP-dependent enzymes that catalyze L-aspartate-dependent decarboxylative Mannich reactions with imine electrophiles have not been identified.

Guided by this UstD-type PLP chemistry, we examined KpbH as a candidate C–C bond-forming enzyme because it shows sequence similarity to UstD (47% identity).^7^ Sequence alignment of KpbH with UstD homologs revealed conservation of Lys232 (Figure S3), which corresponds to the lysine residue predicted to form the internal Schiff base with PLP, supporting the assignment of KpbH as a PLP-dependent enzyme. We therefore hypothesized that KpbH activates L-aspartate as a PLP-bound nucleophilic intermediate and couples it with the iminium carbon of P5C. To test this hypothesis, P5C first had to be generated in situ because this cyclic imine is chemically unstable.

Because the isotope-labeling results implicated an ornithine/P5C-derived precursor for the proline-like unit, we next examined enzymes that could supply P5C under KCP-producing conditions. The *kpb* cluster contains the *kpbG* gene encoding an arginase, and RNA-seq analysis under KCP-producing conditions, achieved by supplementation with L-arginine (RNA-seq raw data: DDBJ/EMBL/NCBI BioProject accession PRJDB37772),^7^ showed upregulation of the *kpbG* gene together with other *kpb* biosynthetic genes. In addition, an ornithine aminotransferase (OAT) gene located outside the *kpb* cluster was also upregulated approximately 4.4-fold under the same conditions. Together, these observations suggest an induced L-arginine-to-L-ornithine-to-P5C route under KCP-producing conditions.

To test whether this OAT could function as a physiological P5C supplier, the endogenous *E. shearii* OAT gene was expressed in *Escherichia coli*, and the recombinant protein, designated EsOAT, was purified. LC–MS analysis confirmed that purified EsOAT indeed generated P5C and L-glutamate from L-ornithine and 2-oxoglutarate (Figures S4 and S5). However, the low expression level and purification yield of EsOAT limited its use in preparative coupled assays and isotope-labeling experiments. We therefore used the *Bacillus subtilis* ornithine aminotransferase BsOAT, also known as RocD,^19,20^ as a more robust in situ P5C-generating enzyme for subsequent biochemical analysis (Figure S4). Because BsOAT generated P5C under the assay conditions, as did EsOAT, it was suitable for evaluating the KpbH-catalyzed reaction.

When P5C was generated in situ in the presence of L-aspartate and KpbH, a new product was detected as an [M+H]^+^ ion at *m/z* 203.1026 (Figure 2). The complete coupled reaction containing L-ornithine, 2-oxoglutarate, L-aspartate, BsOAT, and KpbH gave a clear LC–MS peak corresponding to this product. Product formation depended on both enzymes and all reaction components: the peak was abolished by omission of L-ornithine, L-aspartate, or 2-oxoglutarate, as well as by heat inactivation of BsOAT and/or KpbH. These results indicate that KpbH couples L-aspartate with P5C generated in situ by BsOAT to form the product.

**Figure 2.**
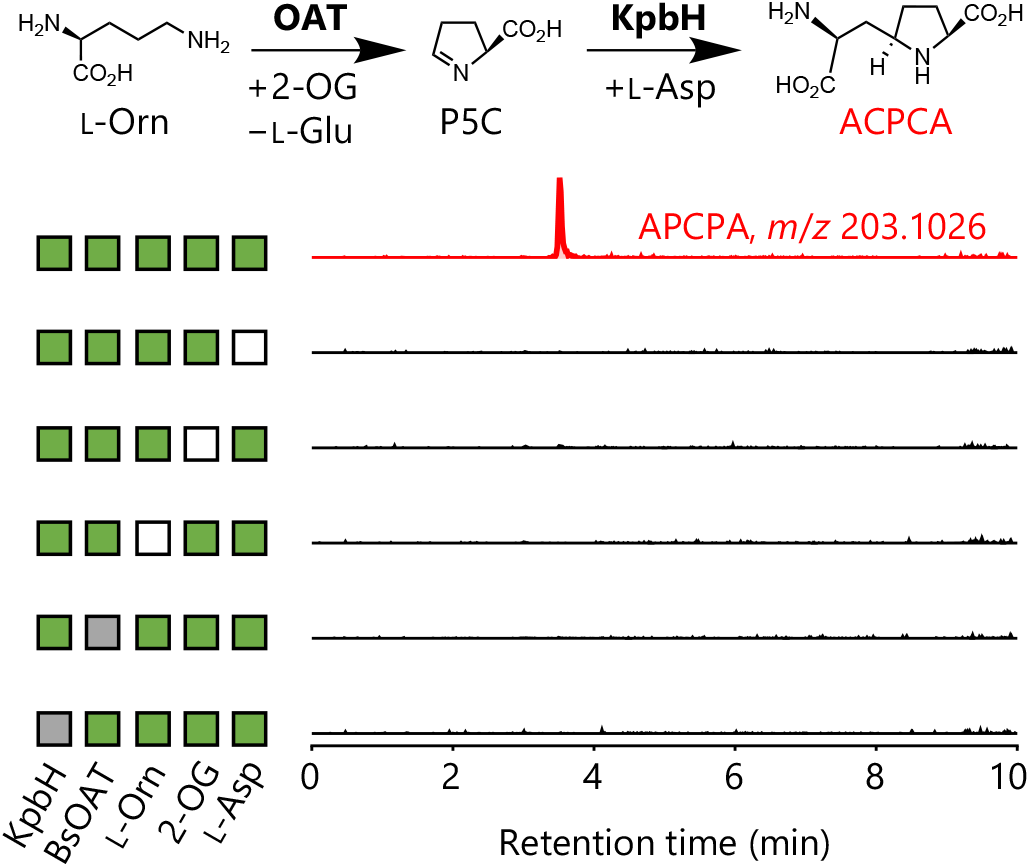
LC-MS analysis of ACPCA formation by the coupled BsOAT/KpbH reaction. EICs for protonated ACPCA ([M+H]^+^, *m/z* 203.1026) are shown. ACPCA was detected only in the complete reaction containing L-ornithine (L-Orn), 2-oxoglutarate (2-OG), L-aspartate, BsOAT, and KpbH. Product formation was abolished by omission of L-aspartate, 2-OG, or L-ornithine (white), or by heat inactivation of BsOAT or KpbH (gray).

The enzymatic product was isolated and analyzed by HRMS and extensive 1D and 2D NMR spectroscopy, establishing its planar structure as 5-(2-amino-2-carboxyethyl)pyrrolidine-2-carboxylic acid, hereafter referred to as ACPCA (Figure S6). Diagnostic NMR correlations established that ACPCA is a C– C-coupled adduct in which the L-aspartate-derived unit is connected to the P5C-derived pyrrolidine-2-carboxylic acid moiety. The C2 stereocenter of the pyrrolidine-2-carboxylic acid moiety inherited from L-ornithine/P5C was assigned as *S* based on its biosynthetic origin, and a diagnostic NOE correlation between H2 and H5 established the relative configuration of the newly formed stereocenter in the pyrrolidine ring, allowing assignment of the ring configuration as 2*S*,5*R*. The *S* configuration at C2 of the 2-amino-2-carboxyethyl substituent, corresponding to C7 in the ACPCA numbering scheme, was independently determined by ^3^*J*_C–H_ and ^3^*J*_H–H_ based configurational analysis (Figures S6A and B).^21,22^ Together, these data established ACPCA as (2*S*,5*R*)-5-((*S*)-2-amino-2-carboxyethyl)pyrrolidine-2-carboxylic acid.

To gain mechanistic insight into the KpbH-catalyzed reaction, we performed the coupled enzymatic reaction in D_2_O. LC–MS analysis revealed formation of ACPCA with a mass increase of +1, corresponding to the M+1 product ion (Figure S7). Subsequent purification, LC–MS/MS analysis, and ^1^H NMR analysis showed that the hydrogen atom at C7 of ACPCA was replaced by deuterium (Figures 3 and S8). In contrast, H_2_O-derived ACPCA retained the C7 proton during ^1^H NMR analysis in D_2_O, indicating that facile nonenzymatic H/D exchange at this position did not occur under the analytical conditions. These results indicate that the C7 hydrogen is introduced during the KpbH-catalyzed reaction through a solvent-exchangeable, enzyme-controlled protonation step. The formation of a single major ACPCA stereoisomer, together with C7 deuterium incorporation in D_2_O, suggests that KpbH controls the facial selectivity of this protonation/deuteration step.

**Figure 3.**
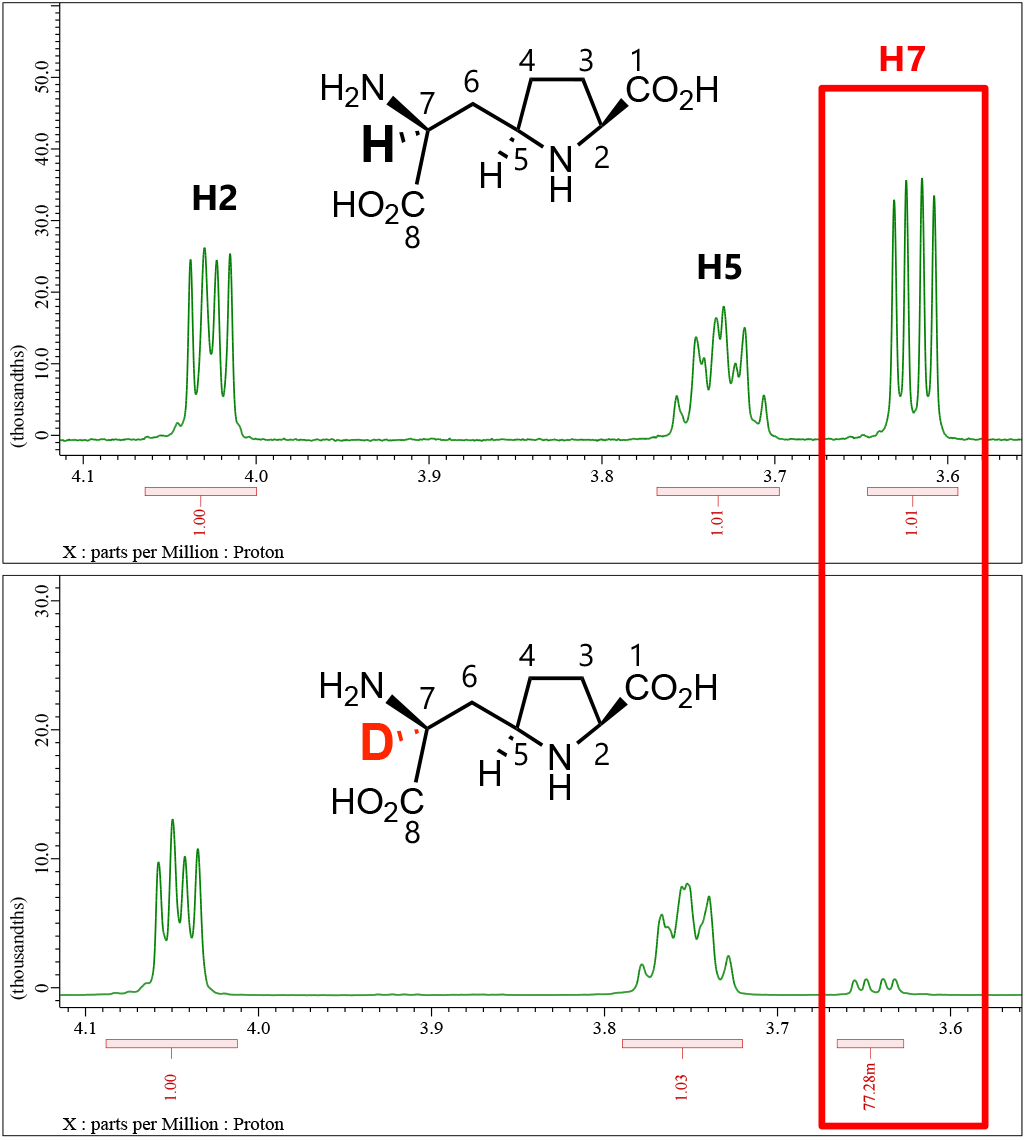
^1^H NMR analysis of deuterium incorporation into ACPCA. Expanded ^1^H NMR spectra of isolated ACPCA generated in H_2_O (top) and D_2_O (bottom) are shown. Both spectra were recorded in D_2_O. In D_2_O-derived ACPCA, the H7 signal was strongly diminished in the D_2_O-derived product, whereas other diagnostic proton signals, including H2 and H5, were retained.

Based on these observations, we propose the following mechanism for the KpbH-catalyzed decarboxylative Mannich-type reaction (Figure 4). In the resting state, KpbH contains PLP as an internal aldimine with Lys232. Upon binding of L-aspartate, transaldimination converts the PLP–Lys232 internal aldimine into the PLP-bound L-aspartate external aldimine. Deprotonation at the Cα-position, followed by decarboxylation of the side-chain carboxylate, generates a PLP-stabilized enamine nucleophile. This intermediate then attacks the iminium carbon of P5C in a Mannich-type C–C bond-forming step. Subsequent enzyme-controlled, solvent-mediated protonation at C7, supported by deuterium incorporation when the reaction is conducted in D_2_O, yields ACPCA and regenerates the PLP cofactor.

**Figure 4.**
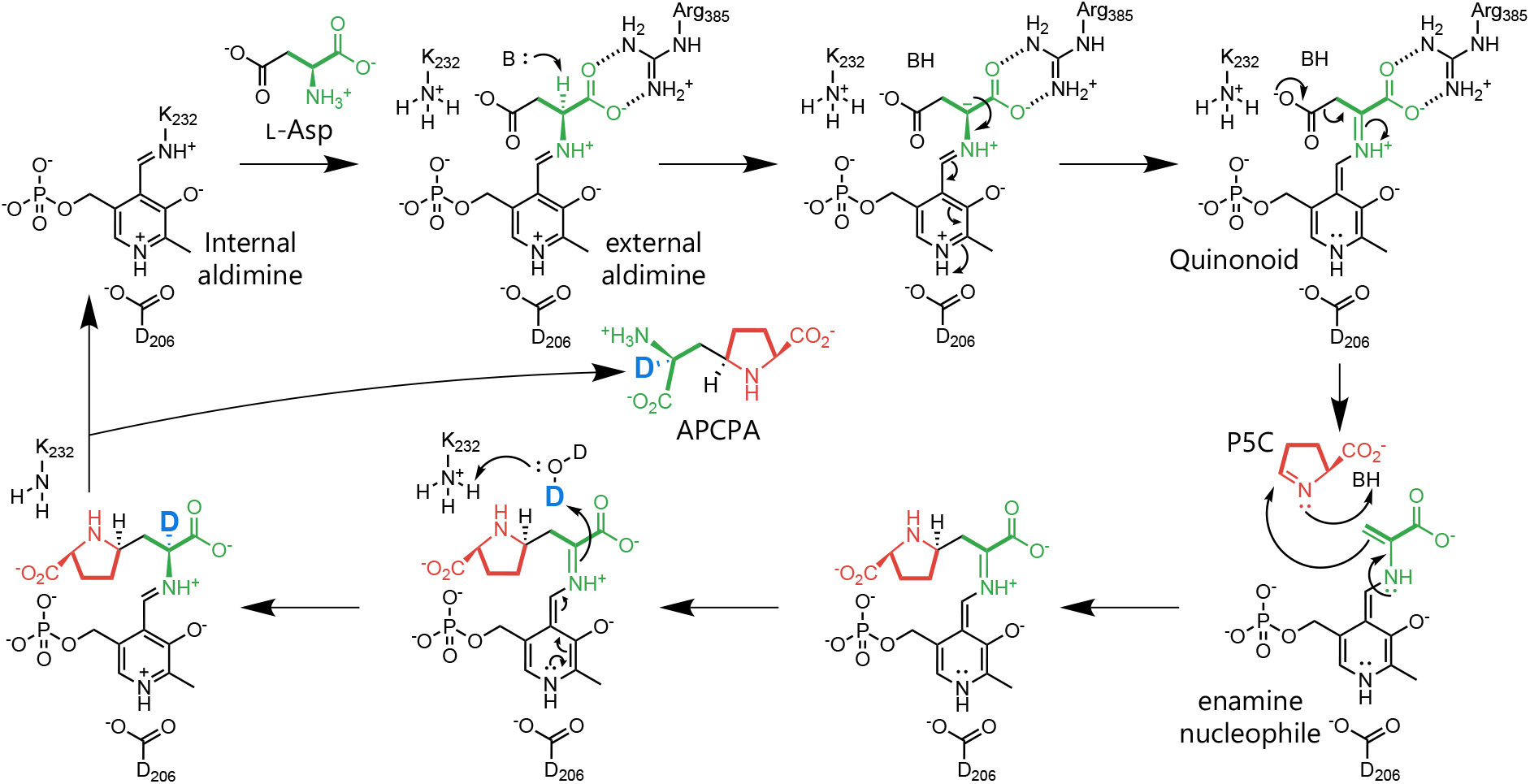
Proposed mechanism of the KpbH-catalyzed PLP-dependent decarboxylative Mannich-type C–C bond-forming reaction. The proposed catalytic sequence is described in the text. Asp206, a conserved acidic residue among PLP-dependent enzymes,^23^ is positioned near the pyridine nitrogen of PLP and is proposed to support the PLP electron sink and stabilize the quinonoid/enamine intermediate. Colored atoms indicate their proposed origins: red, P5C-derived unit; green, L-aspartate-derived unit; blue, solvent-derived deuterium.

AlphaFold3-based complex modeling^24^ further supported this mechanistic proposal (Figure S9). In the modeled PLP–L-aspartate external aldimine complex, Lys232 corresponds to the conserved lysine residue^23,25^ predicted to form the internal Schiff base with PLP in the resting state. Arg385 forms polar contacts with the main-chain carboxylate of L-aspartate and is therefore proposed to anchor the substrate and orient the PLP–L-aspartate external aldimine in a stereoelectronically favorable conformation for quinonoid/enamine formation. Although the catalytic base responsible for α-proton abstraction remains unidentified, the modeled active-site arrangement would align the α-C–H bond of L-aspartate with the PLP π-system in a manner consistent with Dunathan’s stereoelectronic hypothesis.^26^ Thus, the KpbH active site appears to be organized to facilitate formation of the PLP-stabilized enamine intermediate from L-aspartate.

We next examined whether ACPCA is connected to KCP biosynthesis in *E. shearii*. An ACPCA-containing BsOAT/KpbH reaction mixture prepared in D_2_O was lyophilized, redissolved in H_2_O, and applied to *E. shearii* culture plates. Feeding of this D_2_O-prepared reaction mixture increased the M+1/M+0 peak-area ratio of protonated KCP ions to 26.8%, exceeding the calculated natural-abundance M+1/M+0 ratio of 19.5% (Figure S10). This enrichment was not observed to the same extent when the H_2_O-prepared reaction mixture was fed. This selective increase in the M+1/M+0 ratio is consistent with incorporation of deuterium-enriched ACPCA-derived material into KCP.

MS/MS analysis further supported retention of deuterium within the ACPCA-derived portion of KCP, strengthening the connection between ACPCA and KCP biosynthesis (Figure S11). The H_2_O-prepared reaction mixture also increased the total KCP signal (Figure S12), indicating ACPCA-containing material stimulates KCP production. Thus, the moderate deuterium enrichment observed upon feeding of the D_2_O-prepared reaction mixture is consistent with incorporation of ACPCA-derived deuterium into KCP, although concurrent de novo biosynthesis of unlabeled KCP likely diluted the observed enrichment. Together, these feeding experiments support ACPCA as a biosynthetic intermediate in KCP biosynthesis.

In summary, we established that the first C–C bond-forming step in KCP biosynthesis is a stereoselective PLP-dependent decarboxylative Mannich-type reaction catalyzed by KpbH, in which L-aspartate and P5C are coupled to generate ACPCA. This finding provides direct biochemical support for dual utilization of L-ornithine in KCP biosynthesis (Figure 1). In this model, EsOAT converts the carbon skeleton of L-ornithine into P5C for incorporation into the proline-derived unit, whereas KpbB uses the terminal amino group of L-ornithine as the amino donor in the late-stage transamination step. Thus, L-ornithine is utilized in two chemically distinct forms during KCP biosynthesis.

Although native PLP-dependent Mannich enzymes such as LolT have been reported (Figure S13),^27^ native PLP-dependent decarboxylative Mannich-type reactions remain largely unexplored in natural product biosynthesis. KpbH therefore represents the first native PLP-dependent enzyme shown to catalyze an L-aspartate-dependent decarboxylative Mannich-type C–C bond-forming reaction in natural product biosynthesis. More broadly, KpbH demonstrates how an L-aspartate-derived nucleophile can be merged with a cyclic imine electrophile to construct a densely functionalized, noncanonical amino acid-like scaffold in a native biosynthetic pathway. Further exploration and engineering of KpbH may provide new biocatalysts for accessing noncanonical proline-derived amino acid scaffolds.

## Author Contributions

All authors have given approval to the final version of the manuscript.

## Notes

The authors declare no competing financial interest.

## ACKNOWLEDGEMENTS

The authors are grateful to Hiroyuki Watanabe and Kazuo Furihata (The University of Tokyo) for their kind assistance with the NMR measurements. This work was supported by JSPS KAKENHI Grant Number 22H05120 to T.K.

## REFERENCES

(1) Shin-ya, K.; Kim, J.-S.; Furihata, K.; Hayakawa, Y.; Seto, H. Structure of Kaitocephalin, a Novel Glutamate Receptor Antagonist Produced by Eupenicillium Shearii. Tetrahedron Lett. 1997, 38 (40), 7079–7082.

(2) Limon, A.; Reyes-Ruiz, J. M.; Vaswani, R. G.; Chamberlin, A. R.; Miledi, R. Kaitocephalin Antagonism of Glutamate Receptors Expressed in Xenopus Oocytes. ACS Chem. Neurosci. 2010, 1 (3), 175–181.

(3) Zimmer, L.; Crüsemann, M. Amino Acid Tailoring Strategies in Peptide Natural Product Biosynthesis. Trends Chem. 2025, 7 (11), 705–718.

(4) Süssmuth, R. D.; Mainz, A. Nonribosomal Peptide Synthesis-Principles and Prospects. Angew. Chem. Int. Ed Engl. 2017, 56 (14), 3770–3821.

(5) Grininger, M. Enzymology of Assembly Line Synthesis by Modular Polyketide Synthases. Nat. Chem. Biol. 2023, 1–15.

(6) Whitehead, J. N.; Leferink, N. G. H.; Johannissen, L. O.; Hay, S.; Scrutton, N. S. Decoding Catalysis by Terpene Synthases. ACS Catal. 2023, 13 (19), 12774–12802.

(7) Maeno, Y.; Shiraishi, T.; Saito, N.; Maruyama, J.-I.; Shin-Ya, K.; Kuzuyama, T. Biosynthesis of Kaitocephalin: A Neuroprotective Natural Product Featuring a Peptide-like yet Nonpeptidic Scaffold. Angew. Chem. Int. Ed Engl. 2026, No. e23010, e23010.

(8) Du, Y.-L.; Ryan, K. S. Pyridoxal Phosphate-Dependent Reactions in the Biosynthesis of Natural Products. Nat. Prod. Rep. 2019, 36 (3), 430–457.

(9) Daniel-Ivad, P.; Ryan, K. S. New Reactions by Pyridoxal Phosphate-Dependent Enzymes. Curr. Opin. Chem. Biol. 2024, 81, 102472.

(10) Mizutani, T.; Abe, I. Expanding the Catalytic Repertoire of C-C Bond-Forming Pyridoxal 5’-Phosphate-Dependent Enzymes for Noncanonical Amino Acid Formation. Curr. Opin. Biotechnol. 2026, 97 (103437), 103437.

(11) Ye, Y.; Minami, A.; Igarashi, Y.; Izumikawa, M.; Umemura, M.; Nagano, N.; Machida, M.; Kawahara, T.; Shin-Ya, K.; Gomi, K.; Oikawa, H. Unveiling the Biosynthetic Pathway of the Ribosomally Synthesized and Post-Translationally Modified Peptide Ustiloxin B in Filamentous Fungi. Angew. Chem. Int. Ed Engl. 2016, 55 (28), 8072–8075.

(12) Zhang, R.; Tan, J.; Luo, Z.; Dong, H.; Ma, N.; Liao, C. Stereo-Selective Synthesis of Non-Canonical γ-Hydroxy-α-Amino Acids by Enzymatic Carbon–Carbon Bond Formation. Catal. Sci. Technol. 2021, 11 (22), 7380–7385.

(13) Ellis, J. M.; Campbell, M. E.; Kumar, P.; Geunes, E. P.; Bingman, C. A.; Buller, A. R. Biocatalytic Synthesis of Non-Standard Amino Acids by a Decarboxylative Aldol Reaction. Nat. Catal. 2022, 5 (2), 136–143.

(14) Lan, Y.; Zhang, C.; Tang, C.; Ye, Y.; Zhang, R.; Sheng, X.; Liao, C. Semirational Protein Engineering of a Decarboxylative Aldolase for the Regiodivergent and Stereodivergent Synthesis of Cyclic Imino Acids. Angew. Chem. Int. Ed Engl. 2025, 64 (11), e202500080.

(15) Wang, L.; Zhang, D.; Yin, K.; Wu, G.; Duan, X.; Yang, C.; Ma, L.; Chen, X.; Xu, J. Enzymatic Mannich Reaction for the Synthesis of γ-Chiral Amino-Substituted α-Amino Acids. ACS Catal. 2026, No. acscatal.5c07217. 10.1021/acscatal.5c07217.

(16) Lan, Y.; Zhang, L.; Yu, K.; Zhang, R.; Liao, C. Mannich Reaction with Pyridoxal 5’-phosphate Dependent Decarboxylative Aldolase for Synthesis of Noncanonical α-Amino Acids. ChemistryEurope 2026, 4 (1). 10.1002/ceur.202500486.

(17) Campbell, M. E.; Ohler, A. R.; McGill, M. J.; Buller, A. R. Promiscuity Guided Evolution of Decarboxylative Aldolases for Synthesis of Tertiary γ-Hydroxy Amino Acids. Angew. Chem. Int. Ed Engl. 2025, 64 (15), e202422109.

(18) Zhang, Y.; Lan, Y.; Fan, R.; Feng, L.; Wang, G.; Wu, X.; Wen, L.; Duan, Z.; Xia, Y.; Wang, X.; Zhang, L.; Zhou, L.; Tan, M.; Liao, C.; Lu, X. Biocatalytic- and Chemoproteomic-Guided Discovery of a PHGDH Inhibitor from Chemoenzymatic-Promoted DNA-Encoded Libraries Selection Platform. J. Am. Chem. Soc. 2025, 147 (48), 44396–44405.

(19) Gardan, R.; Rapoport, G.; Débarbouillé, M. Expression of the rocDEF Operon Involved in Arginine Catabolism in Bacillus Subtilis. J. Mol. Biol. 1995, 249 (5), 843–856.

(20) Warneke, R.; Garbers, T. B.; Herzberg, C.; Aschenbrandt, G.; Ficner, R.; Stülke, J. Ornithine Is the Central Intermediate in the Arginine Degradative Pathway and Its Regulation in Bacillus Subtilis. J. Biol. Chem. 2023, 299 (7), 104944.

(21) Furihata, K.; Seto, H. J-Resolved HMBC, a New NMR Technique for Measuring Heteronuclear Long-Range Coupling Constants. Tetrahedron Lett. 1999, 40 (34), 6271–6275.

(22) Hansen, P. E. Carbon—Hydrogen Spin—Spin Coupling Constants. Prog. Nucl. Magn. Reson. Spectrosc. 1981, 14 (4), 175–295.

(23) Toney, M. D. Aspartate Aminotransferase: An Old Dog Teaches New Tricks. Archives of Biochemistry and Biophysics. Academic Press February 15, 2014, pp 119–127. 10.1016/j.abb.2013.10.002.

(24) Abramson, J.; Adler, J.; Dunger, J.; Evans, R.; Green, T.; Pritzel, A.; Ronneberger, O.; Willmore, L.; Ballard, A. J.; Bambrick, J.; Bodenstein, S. W.; Evans, D. A.; Hung, C.-C.; O’Neill, M.; Reiman, D.; Tunyasuvunakool, K.; Wu, Z.; Žemgulytė, A.; Arvaniti, E.; Beattie, C.; Bertolli, O.; Bridgland, A.; Cherepanov, A.; Congreve, M.; Cowen-Rivers, A. I.; Cowie, A.; Figurnov, M.; Fuchs, F. B.; Gladman, H.; Jain, R.; Khan, Y. A.; Low, C. M. R.; Perlin, K.; Potapenko, A.; Savy, P.; Singh, S.; Stecula, A.; Thillaisundaram, A.; Tong, C.; Yakneen, S.; Zhong, E. D.; Zielinski, M.; Žídek, A.; Bapst, V.; Kohli, P.; Jaderberg, M.; Hassabis, D.; Jumper, J. M. Accurate Structure Prediction of Biomolecular Interactions with AlphaFold 3. Nature 2024, 1–3.

(25) Liang, J.; Han, Q.; Tan, Y.; Ding, H.; Li, J. Current Advances on Structure-Function Relationships of Pyridoxal 5’-Phosphate-Dependent Enzymes. Frontiers in Molecular Biosciences. Frontiers Media S.A. March 5, 2019, p 4. 10.3389/fmolb.2019.00004.

(26) Dunathan, H. C. Conformation and Reaction Specificity in Pyridoxal Phosphate Enzymes. Proceedings of the National Academy of Sciences 1966, 55 (4), 712–716.

(27) Gao, J.; Liu, S.; Zhou, C.; Lara, D.; Zou, Y.; Hai, Y. A Pyridoxal 5’-Phosphate-Dependent Mannich Cyclase. Nat. Catal. 2023, 6 (6), 476–486.

