## Supporting Information for "A PLP-Dependent Decarboxylative Mannich Reaction Initiates Construction of the Nonpeptidic Scaffold of Kaitocephalin"

**Experimental Section** 3
General 3
Strains and vectors 3
Cloning 3
Comparative transcriptome analysis 3
Purification of recombinant proteins from *E. coli* BL21(DE3) 4
In vitro assay for the EsOAT and BsOAT reaction. 4
In vitro assay for the BsOAT and KpbH reaction. 5
Purification of ACPCA 5
Kaitocephalin production and tracing experiment with deuterium-labeled ACPCA using *E. shearii* 5

**Supplementary Table** 6
Table S1. List of primers used in this study. 6

**Supplementary Figures** 7
Figure S1. Organization and functional annotation of the *kpb* cluster. 7
Figure S2. Native and engineered reactivity of UstD-type PLP-dependent enzymes 8
Figure S3. Multiple sequence alignment of KpbH with UstD homologs and other PLP-dependent enzyme using L-Asp 9
Figure S4. SDS-PAGE analysis of purified recombinant EsOAT, BsOAT, and KpbH 10
Figure S5. LC-MS analysis of P5C and L-glutamate formation by ornithine aminotransferases 11
Figure S6. NMR analysis of ACPCA and comparison with the corresponding partial structure of KCP. 16
Figure S7. $D_2O$-labeling analysis of ACPCA formation by the coupled BsOAT/KpbH reaction. 17
Figure S8. MS/MS analysis of isolated ACPCA and deuterium-labeled ACPCA. 18
Figure S9. AlphaFold3-based model of the KpbH PLP–L-aspartate external aldimine complex. 19
Figure S10. Incorporation of deuterium-enriched ACPCA-derived material into KCP. 20
Figure S11. MS/MS analysis of KCP and its M+1 ion obtained in the feeding experiment. 21
Figure S12. Effect of reaction mixture feeding on KCP production 22
Figure S13. Proposed catalytic mechanism of the PLP-dependent Mannich cyclase LolT. 23

**REFERENCES** 24

### Experimental Section

#### General

Biochemicals and enzymes for genetic manipulation were purchased from TaKaRa Bio (Ohtsu, Japan), New England Biolabs Japan (Tokyo, Japan) and TOYOBO (Osaka, Japan). Eluent additives in liquid chromatography were purchased from Sigma-Aldrich (St. Louis, MO). Oligonucleotides used for genetic manipulation (Table S1) were purchased from Fasmac Co., Ltd. (Kanagawa, Japan). All other reagents were purchased from FUJIFILM Wako Pure Chemical Industries (Osaka, Japan), Kanto Chemicals (Tokyo, Japan), Tokyo Chemical Industry (Tokyo, Japan) and Nacalai Tesque (Kyoto, Japan) unless otherwise noted. Cells were disrupted using a Branson Sonifier 250 (Emerson Japan, Tokyo, Japan). UHPLC–MS analysis was performed with a Nexera X3 (Shimadzu, Kyoto, Japan) coupled to an X500R QTOF system (SCIEX, Framingham, MA, USA), which was equipped with an electrospray source operating in the positive-ionization mode.

For HPLC-UV/VIS analysis, X-LC (JASCO, Tokyo) system was used. For HPLC-UV/VIS-MS analysis, ACQUITY UPLC H-Class system/ACQUITY QDa (Waters, Tokyo) was used. DNA manipulation was performed according to the manufacturer's instructions. NMR spectra except for NOESY and *J*-resolved HMBC were obtained using a JEOL ECA-600. NOESY and *J*-resolved HMBC spectra were obtained using a Varian Unity INOVA 500.

#### Strains and vectors

*Escherichia coli* DH5α was used for cloning following standard recombinant DNA techniques. *E. coli* BL21(DE3) was used for expression of protein. *Eupenicillium shearii* PF11911 was used for labeling experiments of KCP and RNA sequencing. *Bacillus subtilis* 168 was used for cloning of BsOAT. pHis8[1] was used for protein expression.

#### Cloning

Plasmid constructs containing codon-optimized synthetic genes encoding KpbH and EsOAT for expression in *E. coli*, cloned into pET-28a(+) to introduce an N-terminal His6 tag, were purchased from Twist Bioscience (San Francisco, CA, USA). The *BsOAT* gene was amplified from *B. subtilis* 168 genomic DNA using the primers listed in Table S1 and inserted into the pHis8 vector linearized with HindIII and NcoI using NEBuilder® HiFi DNA Assembly (New England Biolabs Japan, Tokyo, Japan), yielding a construct encoding N-terminally His8-tagged BsOAT.

#### Comparative transcriptome analysis

Using RNA-seq raw data (DDBJ/EMBL/NCBI BioProject accession PRJDB37772) obtained as previously described in Ref.[2], a de novo transcriptome analysis was performed with OmicsBox to compare a KCP-producing condition, in which mycelia were cultured for 5 days on potato dextrose agar supplemented with adenine and L-arginine, and a non-producing condition cultured for 5 days on potato dextrose agar without supplementation. Gene expression levels were evaluated based on RPKM values. Ornithine aminotransferase located outside the *kpb* cluster (EsOAT) was identified from the *E. shearii* genome through homology search using KpbB as a query. It showed higher expression under the KCP-producing condition, with RPKM values of 4496.734 and 1025.673 under the producing and non-producing conditions, respectively.

#### Purification of recombinant proteins from *E. coli* BL21(DE3)

After each plasmid was introduced into *E. coli* BL21(DE3), the resulting transformants were cultivated in 200 mL of TB medium containing tryptone (1.2%), yeast extract (2.4%), glycerol (0.4%), $KH_2PO_4$ (0.23%), and $K_2HPO_4$ (1.25%), supplemented with kanamycin, at 37 °C for 5 h until the cells reached an $OD_{600}$ of approximately 0.6. After the cultures were cooled on ice for 10 min, gene expression was induced by the addition of isopropyl β-D-1-thiogalactopyranoside (IPTG) at a final concentration of 100 μM, and the cultures were further incubated at 18 °C for 14-18 h. The cells were harvested by centrifugation.

Three recombinant proteins, EsOAT, BsOAT, and KpbH, were purified by Ni-affinity chromatography. All buffers used for purification were supplemented with 50 μM pyridoxal 5′-phosphate (PLP). The harvested cells were resuspended in 20 mL of wash buffer consisting of 50 mM Tris-HCl (pH 8.0), 500 mM NaCl, 20 mM imidazole, 20% glycerol, and 50 μM PLP, and lysed by sonication on ice. Cell debris was removed by centrifugation at 4 °C (30,000 × g, 20 min). The supernatant was loaded onto 1 mL of Ni-NTA agarose resin (FUJIFILM Wako Pure Chemical Corporation, Osaka, Japan), and the resin was washed with 20 mL of wash buffer. The bound proteins were eluted with 2.5 mL of elution buffer consisting of 50 mM Tris-HCl (pH 8.0), 500 mM NaCl, 250 mM imidazole, 20% glycerol, and 50 μM PLP. The eluate was loaded onto a PD-10 column and eluted with 3.5 mL of 100 mM ammonium bicarbonate buffer (pH 7.7). Each purified protein was analyzed by SDS–PAGE. When necessary, the protein solutions were concentrated using a Vivaspin 20 ultrafiltration unit (Sartorius, Göttingen, Germany), flash-frozen in liquid nitrogen, and stored in liquid nitrogen.

#### In vitro assay for the EsOAT and BsOAT reaction.

The standard reaction was performed at 30 °C for 1h in 100 μL reaction mixture containing 100 mM ammonium bicarbonate (pH 7.7), 1 mM L-ornithine, 2 mM 2-oxoglutarate, 50 μM PLP, 5 μM EsOAT, or 5 μM BsOAT. After centrifugation, the supernatant was subjected to LC-UV/VIS-MS analysis. After the reaction, the mixture was quenched by adding an equal volume of acetonitrile. The sample was centrifuged, and the resulting supernatant was subjected to HR-LC-MS/MS analysis.

#### In vitro assay for the BsOAT and KpbH reaction.

The standard reaction was performed at 30 °C for 1h in 100 µL reaction mixture containing 100 mM ammonium bicarbonate (pH 7.7), 1 mM L-ornithine, 2 mM 2-oxoglutarate, 50 µM PLP, 2 mM L-aspartate, 5 µM EsOAT, and 2 µM KpbH. After the reaction, the mixture was quenched by adding an equal volume of acetonitrile. The sample was centrifuged, and the resulting supernatant was subjected to HR-LC-MS/MS analysis.

The reaction mixture 1 for the following tracing experiment containing 400 mM ammonium bicarbonate pH7.7, 40 mM L-ornithine, 80 mM 2-oxoglutarate, 500 µM PLP, 80 mM L-aspartate, 141 µM EsOAT, and 63 µM KpbH, with either $H_2O$ or $D_2O$ as the solvent. An additional reaction was performed under the $D_2O$ condition in the absence of L-ornithine.

#### Purification of ACPCA

ACPCA (**4**) was prepared by enzymatic reaction on a 40 mL scale. The reaction mixture contained 400 mM ammonium bicarbonate (pH 7.7), 40 mM L-ornithine, 80 mM 2-oxoglutarate, 500 µM PLP, 80 mM L-aspartate, 17 µM EsOAT, and 5 µM KpbH. The reaction was performed at 30 °C for 6 h. After incubation, the reaction mixture was lyophilized and redissolved in 5 mL of 50% acetonitrile. Insoluble material was removed by centrifugation (7,030 × g, 10 min), and the resulting supernatant was filtered. The filtrate was fractionated in 300 µL portions using an XBridge BEH Amide Column (130 Å, 5 µm, 4.6 × 250 mm). The mobile phase consisted of 75% acetonitrile containing 10 mM ammonium acetate (pH 3.0). After purification, the collected sample was stored at 4 °C for 1 day, and the resulting precipitate was collected by centrifugation (21,500 × g, 10 min). The collected white solid was lyophilized several times to afford ACPCA (**4**) as a white powder (1.3 mg).

Deuterium-labeled ACPCA was purified under the same conditions to afford a white powder (1.9 mg).

#### Kaitocephalin production and tracing experiment with deuterium-labeled ACPCA using *E. shearii*

Kaitocephalin was produced on plates for 9 days under the same conditions as previously reported,[2] except that 20 mL of potato dextrose agar (PDA) medium were used instead of 130 mL. For the feeding experiments, the reaction mixtures used for ACPCA labeling were prepared on a 10 mL scale and incubated for 3 h. The mixtures were then lyophilized, redissolved in 10 mL of water, and filter sterilized. The resulting solution (1 mL) was spread onto each plate. The harvested mycelia were frozen in liquid nitrogen, ground with a mortar and pestle, extracted with methanol, filtered, and subjected to HR-LC-MS/MS analysis.

#### Supplementary Table

**Table S1. List of primers used in this study.**

| | |
|---|---|
| pHis8-BsOAT_fw | 5′-gtctggttccgcgtggttccACAGCTTTATCTAAATCCAAAGAAATTATTG-3′ |
| pHis8-BsOAT_rv | 5′-ggtgctcgagtgcggccgcaTTATGCGTTTCGCAGCAC-3′ |

#### Supplementary Figures

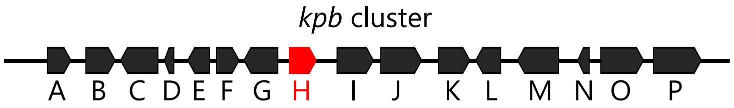


| | |
|---|---|
| KpbA | 4-Hydroxyphenylpyruvate dioxygenase |
| KpbB | Ornithine aminotransferase |
| KpbC | AMP-binding protein |
| KpbD | Acyl carrier protein |
| KpbE | Aldolase |
| KpbF | ACC-synthase |
| KpbG | Arginase |
| KpbH | PLP-dependent enzyme |
| KpbI | 2-OG-dependent dioxygenase |
| KpbJ | FMN-dependent dehydrogenase |
| KpbK | Dipeptidase |
| KpbL | Hypothetical protein |
| KpbM | Flavin-dependent halogenase |
| KpbN | Glutathione *S*-transferase |
| KpbO | MFS transporter |
| KpbP | γ-Glutamyltransferase |

**Figure S1. Organization and functional annotation of the *kpb* cluster.**
The cluster contains 16 open reading frames, *kpbA*–*kpbP*, encoding enzymes and proteins predicted to be involved in kaitocephalin biosynthesis. Predicted functions of the encoded proteins are listed below the gene map. KpbH, a putative PLP-dependent enzyme, was investigated in this study as a PLP-dependent candidate enzyme for C–C bond formation between P5C and L-aspartate.

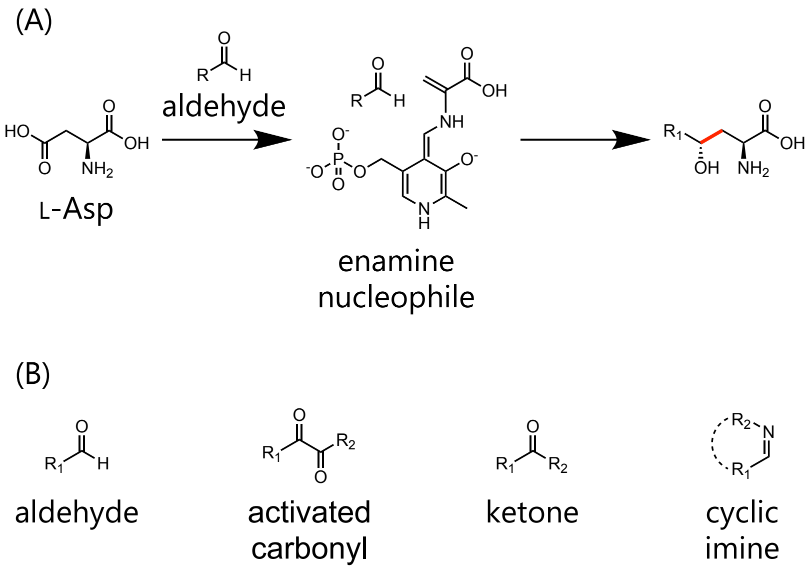


**Figure S2. Native and engineered reactivity of UstD-type PLP-dependent enzymes.** (A) Native UstD catalyzes the decarboxylative aldol-type C–C bond-forming reaction between L-aspartate and an aldehyde electrophile. Decarboxylation of the L-aspartate-derived external aldimine generates a PLP-stabilized enamine nucleophile, which attacks the aldehyde to form a γ-hydroxy amino acid scaffold. The newly formed C–C bond is highlighted in red. (B) Representative classes of electrophiles accepted by engineered UstD variants. UstD-type enzymes have been engineered to react with aldehydes, activated carbonyl compounds, ketones, and cyclic imines, expanding their utility for the synthesis of noncanonical amino acids.

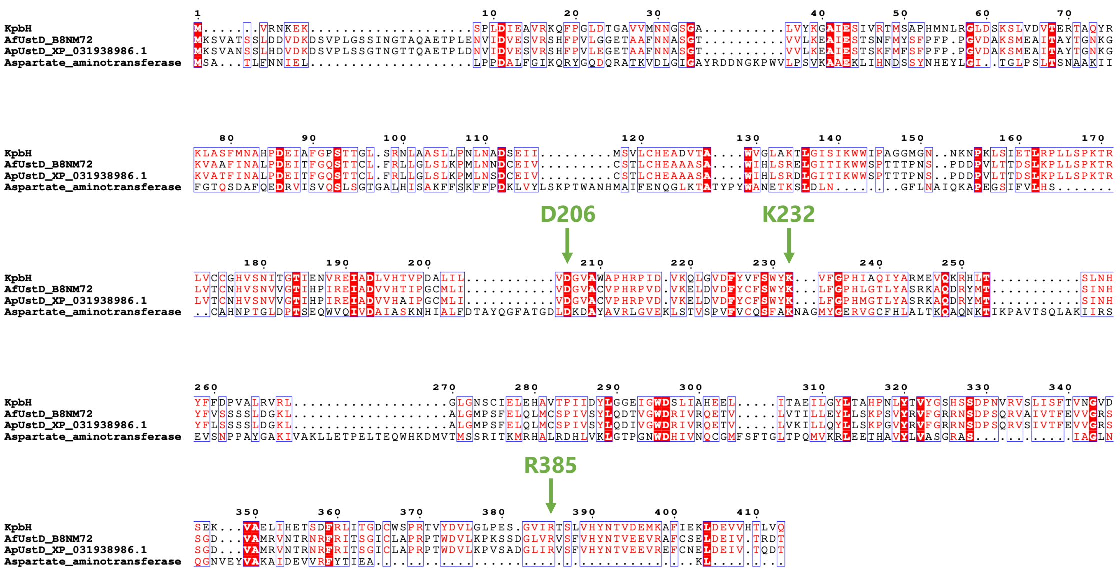

**Figure S3. Multiple sequence alignment of KpbH with UstD homologs and other PLP-dependent enzyme using L-Asp.**

Conserved residues implicated in the proposed reaction mechanism are indicated by green arrows. Asp206 is conserved among the PLP-dependent enzymes analyzed here and is proposed to support the electron-sink function of PLP and stabilize the quinonoid intermediate. Lys232 corresponds to the catalytic lysine that forms the internal aldimine with PLP, whereas Arg385 is conserved among UstD homologs but not in the aminotransferase from *Saccharomyces cerevisiae*. In the modeled PLP- L-aspartate external aldimine complex of KpbH, Arg385 is positioned to interact with the α-carboxylate of L-aspartate. Residue numbering is based on KpbH.

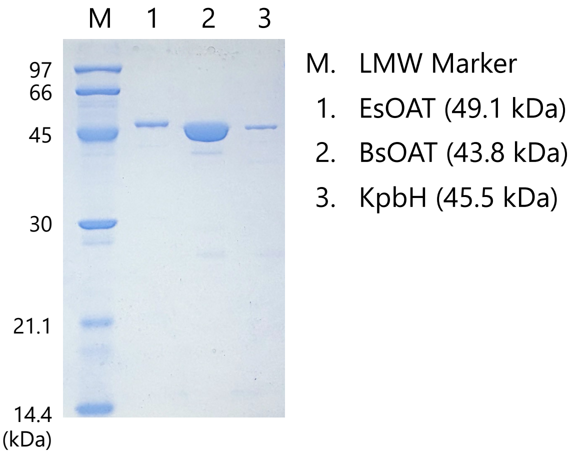


**Figure S4. SDS-PAGE analysis of purified recombinant EsOAT, BsOAT, and KpbH.** Lane M, low-molecular-weight protein marker; lane 1, EsOAT (calculated molecular mass, 49.1 kDa); lane 2, BsOAT/RocD (43.8 kDa); lane 3, KpbH (45.5 kDa). The gel was stained with Coomassie Brilliant Blue.

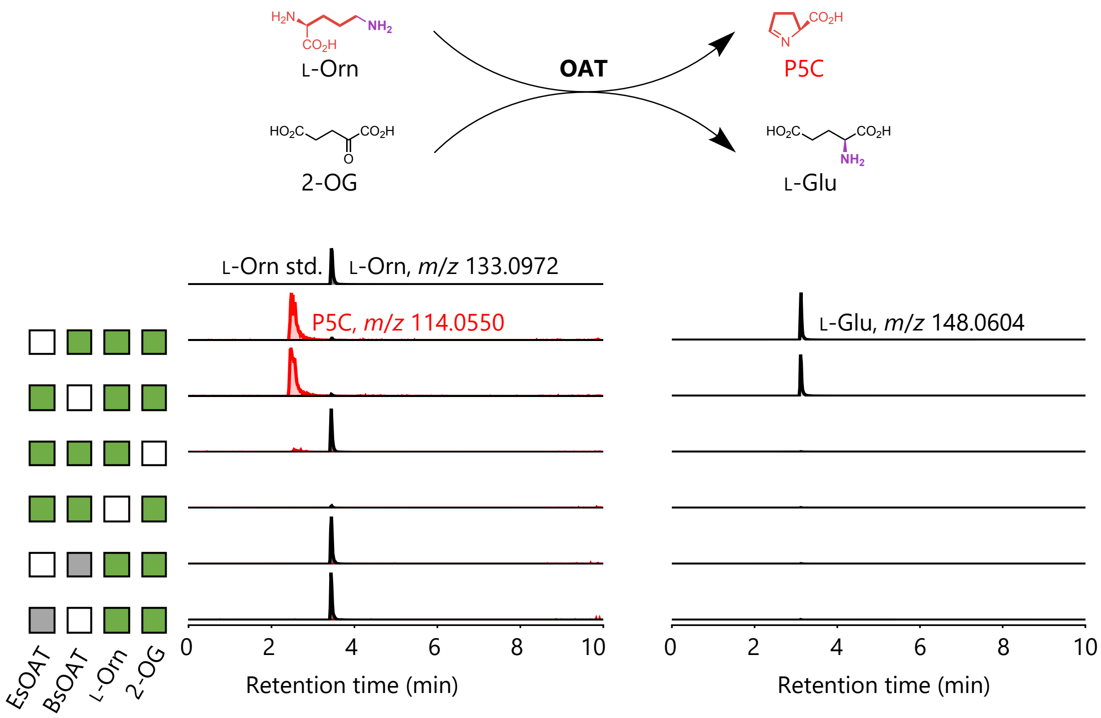


**Figure S5. LC-MS analysis of P5C and L-glutamate formation by ornithine aminotransferases.**

Extracted ion chromatograms (EICs) monitored at *m/z* 133.0972, 114.0550, and 148.0604, corresponding to the protonated molecules ($[M+H]^+$) of L-ornithine (L-Orn), P5C, and L-glutamate (L-Glu), respectively, are shown. In the ornithine aminotransferase reaction, L-Orn is converted to P5C with concomitant formation of L-Glu from 2-oxoglutarate (2-OG). P5C formation was observed in reactions containing L-Orn, 2-OG, and either BsOAT or EsOAT, supporting L-Glu formation through the corresponding transamination reaction. P5C formation was not observed, or was markedly reduced, in the absence of L-Orn, 2-OG, or when heat-inactivated BsOAT or EsOAT was used. These results support the ornithine aminotransferase activity of both enzymes.

(A)

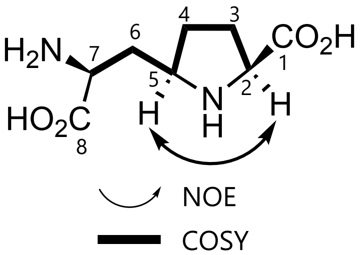


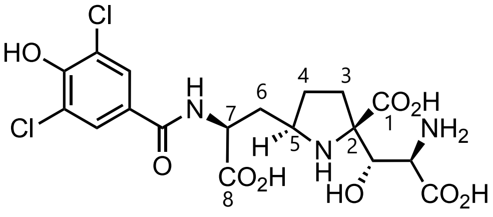


KCP

| No. | $\delta_H$ (ppm) (multi, $J_H$ (Hz)) | δC (ppm) | HMBC correlations |
|---|---|---|---|
| 1 | | 174.6 | |
| 2 | 4.03 (dd, 9.0 4.8) | 61.4 | C1, C3 |
| 3 | Ha: 2.08 (m) | 28.4 | C1, C2, C5 |
| | Hb: 2.17 (m) | | C1, C2, C4 |
| 4 | Ha: 1.58 (m) | 29.4 | C3, C5, C6 |
| | Hb: 2.18 (m) | | C2, C3 |
| 5 | 3.74 (m) | 58.5 | C4, C6, C7 |
| 6 | 2.02 (ddd, 14.4 7.8 4.2) | 33.5 | C4, C5, C7, C8 |
| | 2.29 (ddd, 14.4 9.6 6.6) | | C4, C7, C8 |
| 7 | 3.63 (dd, 9.6 4.2) | 52.4 | C5, C6, C8 |
| 8 | | 173.6 | |

in $D_2O$

| No. | $\delta_H$ (ppm) (multi, $J_H$ (Hz)) | δC (ppm) |
|---|---|---|
| 1 | | 175.0 |
| 2 | | 77.1 |
| 3 | 2.01 (m) | 32.7 |
| | 2.28 (ddd, 14.0 6.0 2.0) | |
| 4 | 1.61 (m) | 30.4 |
| | 2.12 (m) | |
| 5 | 3.70 (m) | 59.7 |
| 6 | 2.06 (m) | 35.6 |
| | 2.41 (ddd, 14.5 7.0 6.0) | |
| 7 | 4.35 (dd, 8.0 6.0) | 54.2 |
| 8 | | 177.8 |

in $D_2O$

(B)

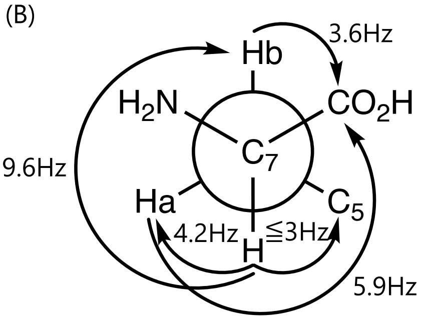

(C) $^1H$

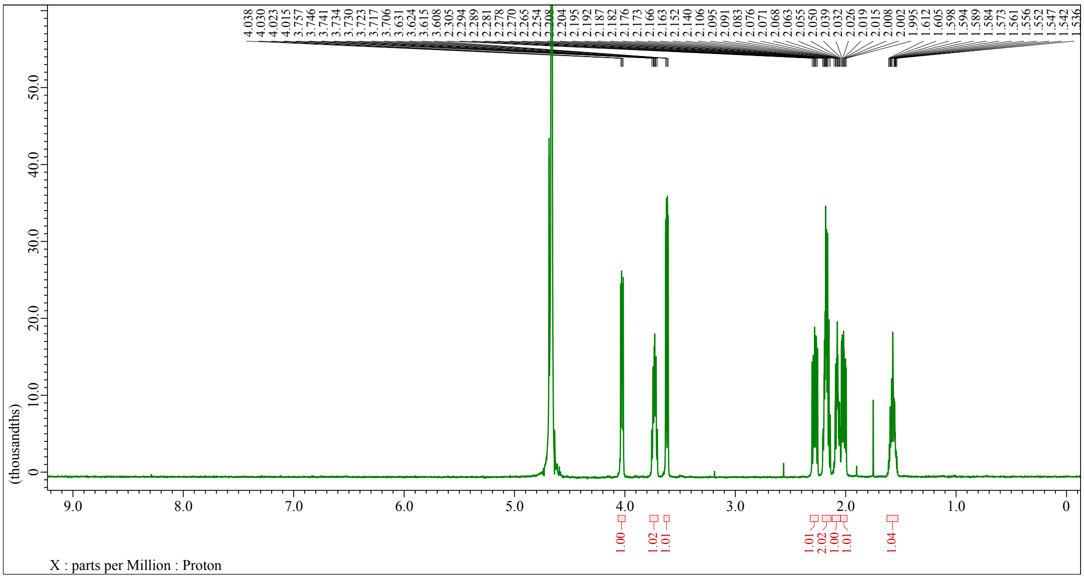

(D) $^{13}C$

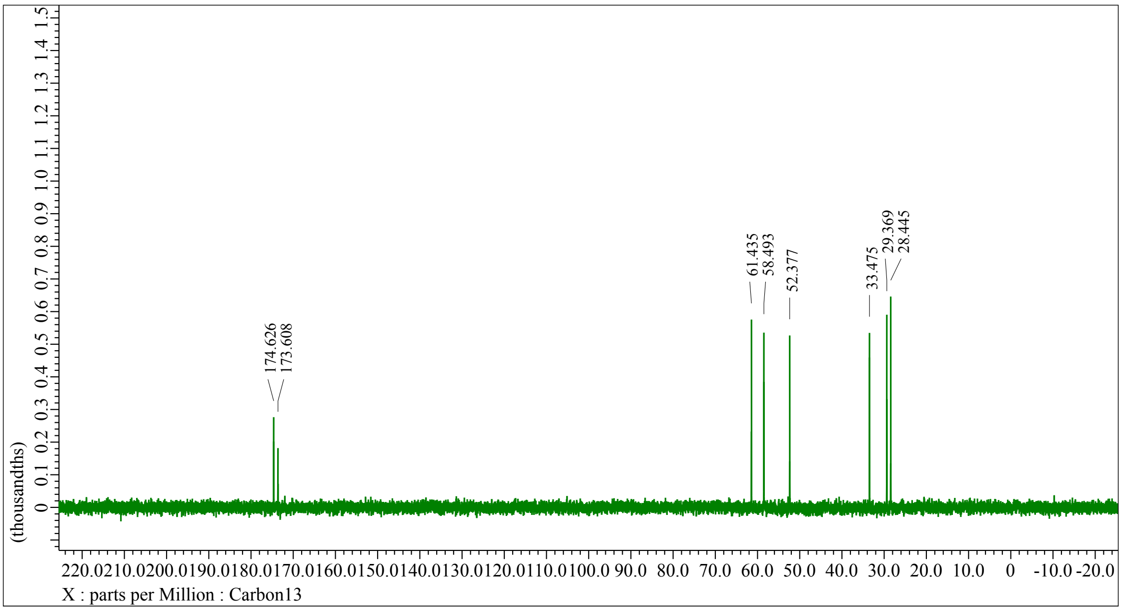

(E) COSY

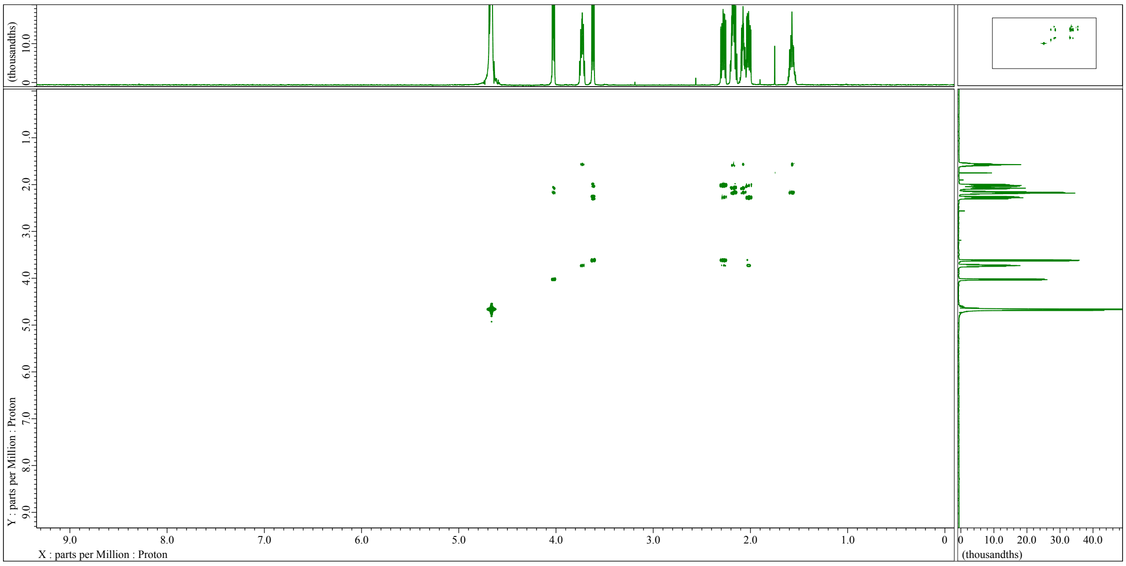
(thousandths)
Y : parts per Million : Proton
X : parts per Million : Proton
(thousandths)

(F) HSQC

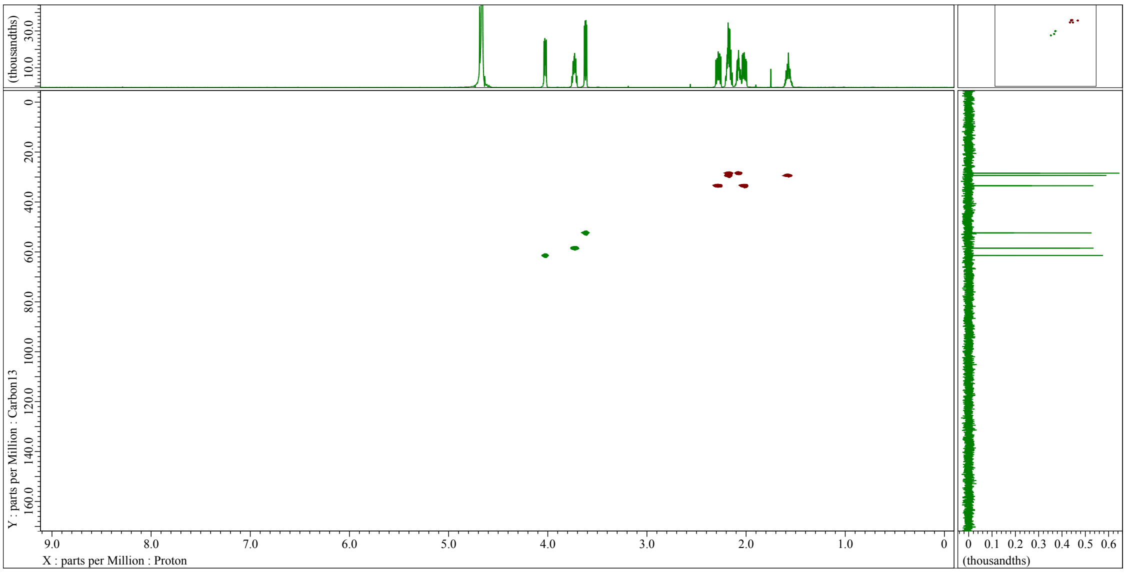
(thousandths)
Y : parts per Million : Carbon13
X : parts per Million : Proton
(thousandths)

(G) HMBC

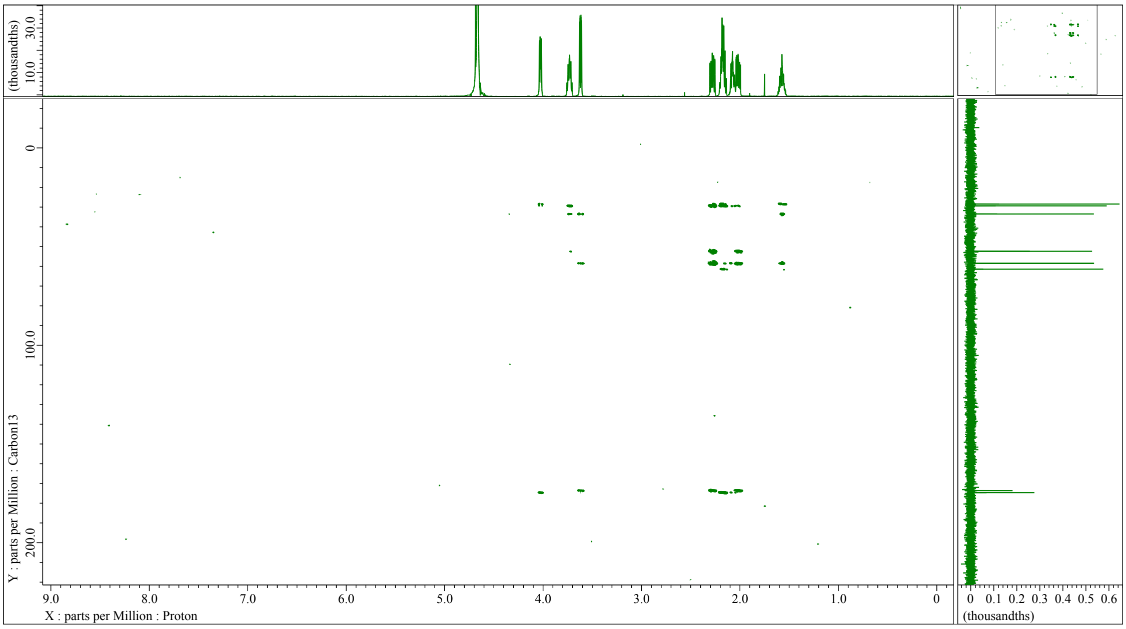


(H) NOESY

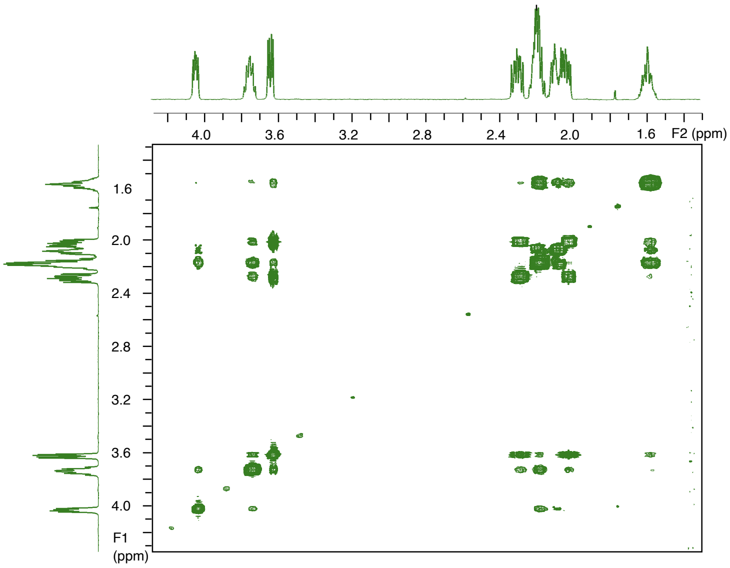

(I)

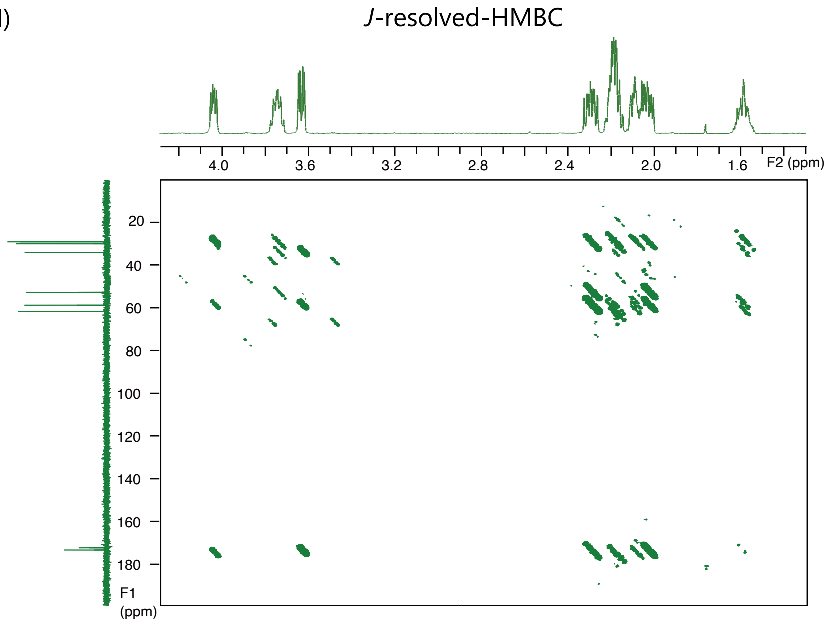


**Figure S6. NMR analysis of ACPCA and comparison with the corresponding partial structure of KCP.**

(A) Key COSY and NOE correlations for ACPCA are shown on the left, and the corresponding partial structure of KCP is shown on the right. NMR assignments in $D_2O$ are summarized in the tables. (B) Fischer projection of the C6–C7 fragment and diagnostic *J* values. The relative configuration of the C6–C7 fragment was assigned by *J*-based configuration analysis using $^{3}J_{H\text{–}H}$ and long-range $^{3}J_{C\text{–}H}$ coupling constants. The observed coupling constants are indicated in Hz. Long-range $^{1}H$–$^{13}C$ coupling constants were obtained by *J*-resolved HMBC according to a previously reported method.[3,4] (C–I) Full $^{1}H$, $^{13}C$, COSY, HSQC, HMBC, NOESY and *J*-resolved HMBC spectra of ACPCA are also included. The NMR data support the assignment of ACPCA as (2*S*,5*R*)-5-((*S*)-2-amino-2-carboxyethyl)pyrrolidine-2-carboxylic acid (ACPCA) and are consistent with the stereochemistry of the corresponding proline-containing partial structure in KCP.

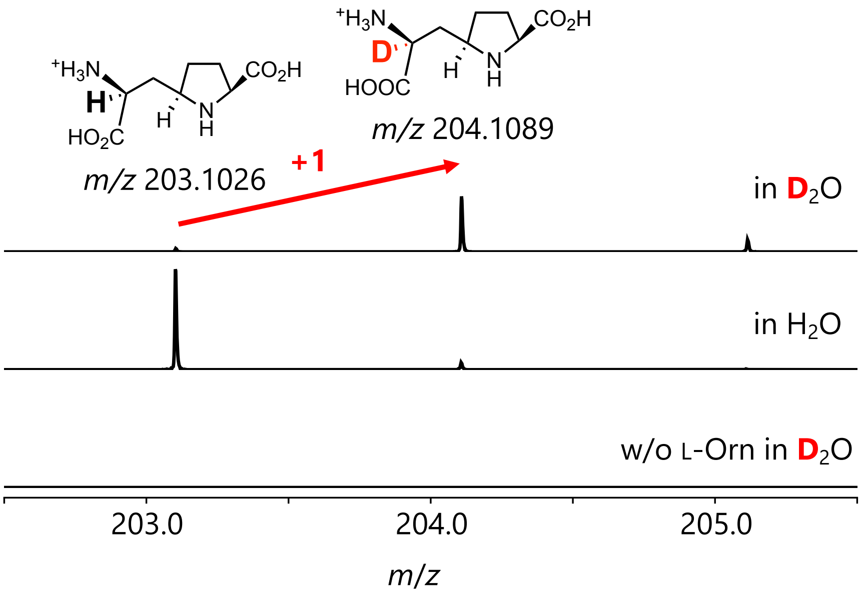


**Figure S7. $D_2O$-labeling analysis of ACPCA formation by the coupled BsOAT/KpbH reaction.**

ACPCA was detected at *m/z* 203.1026 when the reaction was performed in $H_2O$, whereas the reaction in $D_2O$ predominantly gave the M+1 ion at *m/z* 204.1089. Omission of L-ornithine abolished ACPCA formation.

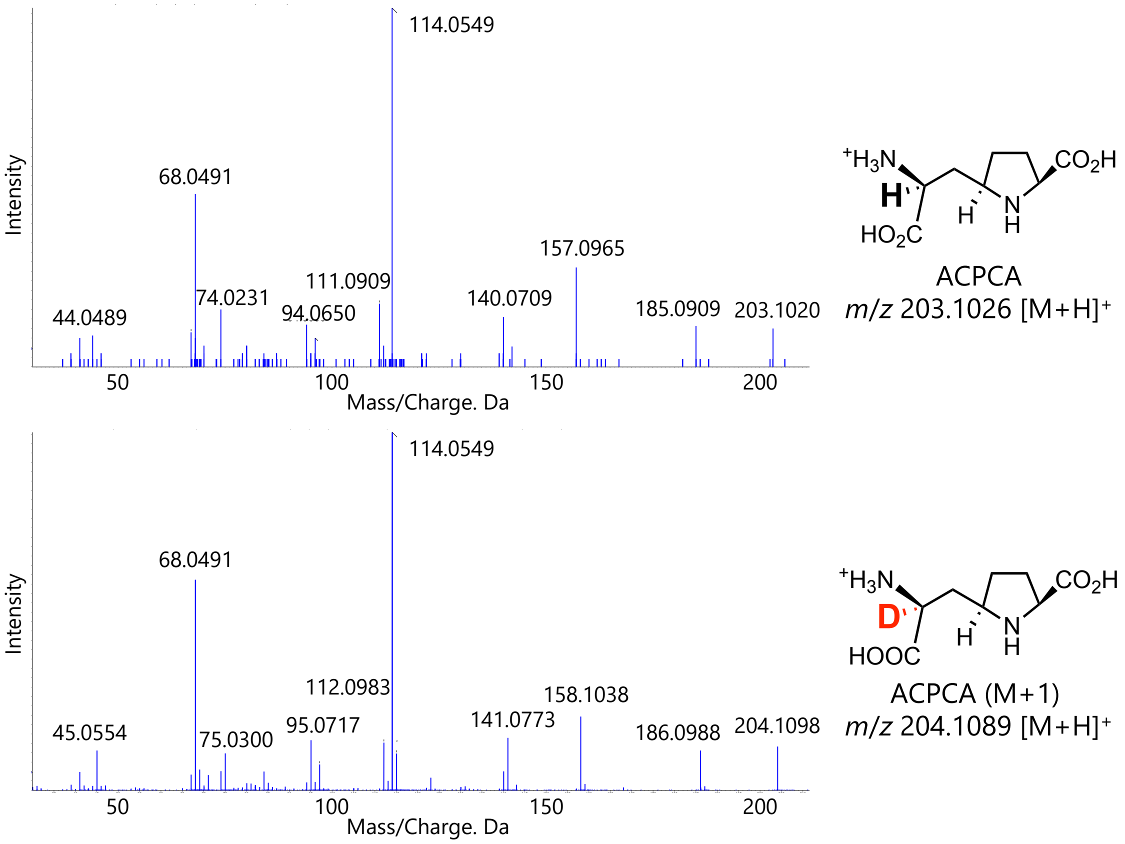


**Figure S8. MS/MS analysis of isolated ACPCA and deuterium-labeled ACPCA.**

MS/MS spectra of isolated ACPCA generated in $H_2O$ (top) and deuterium-labeled ACPCA generated in $D_2O$ (bottom) are shown. The protonated molecular ions were detected at *m/z* 203.1020 and 204.1098, respectively. Selected fragment ions of the deuterium-labeled product showed +1 mass shifts relative to those of unlabeled ACPCA, whereas fragment ions such as *m/z* 114.0549 and 68.0491 remained unchanged. This fragmentation pattern supports deuterium incorporation into the ACPCA scaffold. The position of deuterium incorporation was further assigned by $^1H$ NMR analysis (Figure 3).

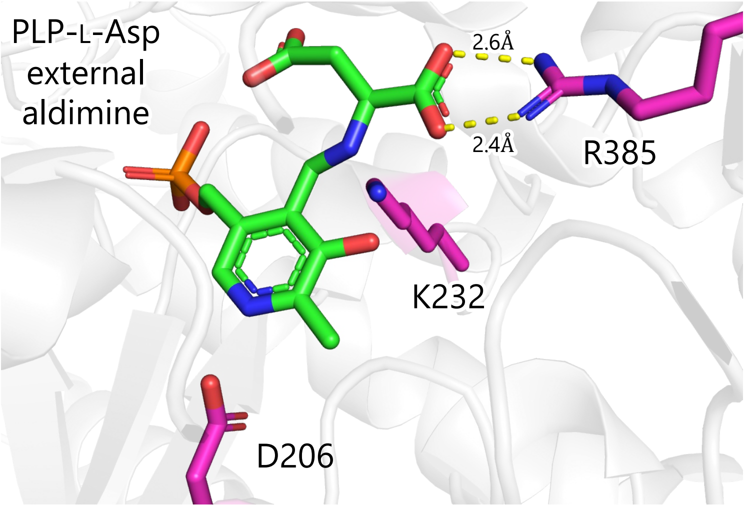


**Figure S9. AlphaFold3-based model of the KpbH PLP–L-aspartate external aldimine complex.**

The modeled PLP– L-aspartate external aldimine is shown as green sticks, and key active-site residues are shown as magenta sticks. Arg385 is positioned to interact with the main-chain carboxylate of L-aspartate through predicted hydrogen-bonding interactions of 2.6 and 2.4 Å, suggesting that this residue helps orient L-aspartate for α-proton abstraction and subsequent decarboxylation. Lys232 corresponds to the catalytic lysine that forms the PLP internal aldimine. Asp206 is located near the PLP pyridinium ring and is proposed to support the PLP electron-sink system, consistent with the conserved role of this acidic residue in stabilizing quinonoid intermediates in PLP-dependent enzymes and UstD homologs.

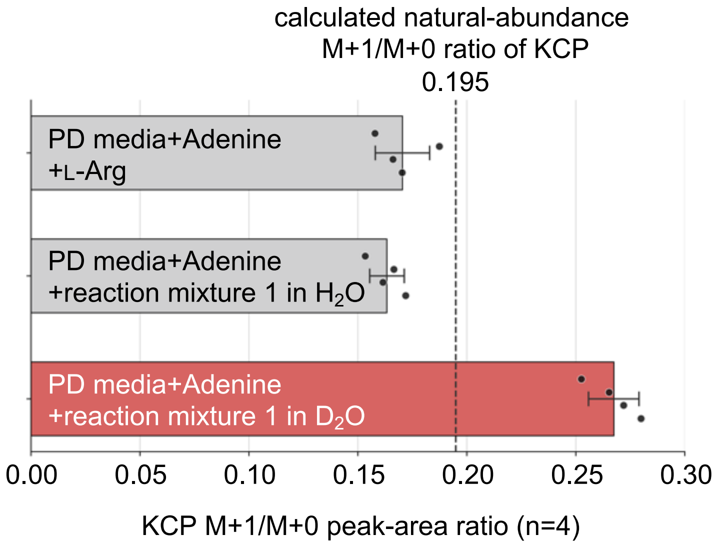


**Figure S10. Incorporation of deuterium-enriched ACPCA-derived material into KCP.** The ACPCA-containing BsOAT/KpbH reaction mixture prepared in $D_2O$ was lyophilized, redissolved in $H_2O$, and applied to *E. shearii* culture plates. The KCP M+1/M+0 peak-area ratio was calculated from peak areas of the protonated monoisotopic KCP ion and the corresponding protonated M+1 isotopologue. Feeding of the $D_2O$-prepared reaction mixture increased the KCP M+1/M+0 ratio above the calculated natural-abundance ratio of 0.195, whereas the $H_2O$-prepared reaction mixture and L-arginine-supplemented control (KCP producing condition in the previous study[2]) remained below this value. Data are shown as mean ± SD from independent biological replicates. Dots indicate individual replicates.

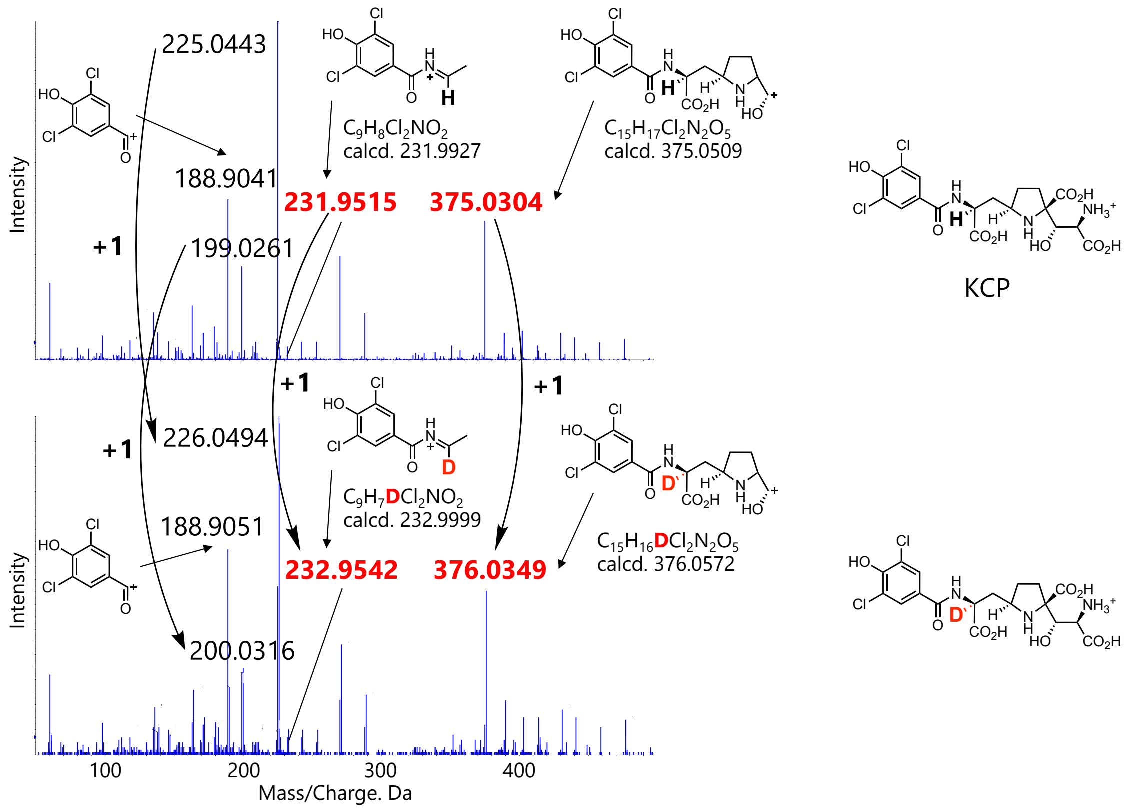


**Figure S11. MS/MS analysis of KCP and its M+1 ion obtained in the feeding experiment.**

MS/MS spectra of unlabeled KCP and the M+1 ion detected after feeding of the $D_2O$-prepared reaction mixture are shown. Fragment ions derived from the ACPCA-derived portion of KCP exhibited corresponding +1 mass shifts in the M+1 ion, including *m/z* 231.9515 to 232.9542 and *m/z* 375.0304 to 376.0349. Additional fragment ions also showed the expected +1 shifts (e.g., *m/z* 225.0443 to 226.0494 and *m/z* 199.0261 to 200.0316). These data support retention of deuterium in the ACPCA-derived moiety of KCP and are consistent with incorporation of deuterium-labeled ACPCA-derived material during KCP biosynthesis.

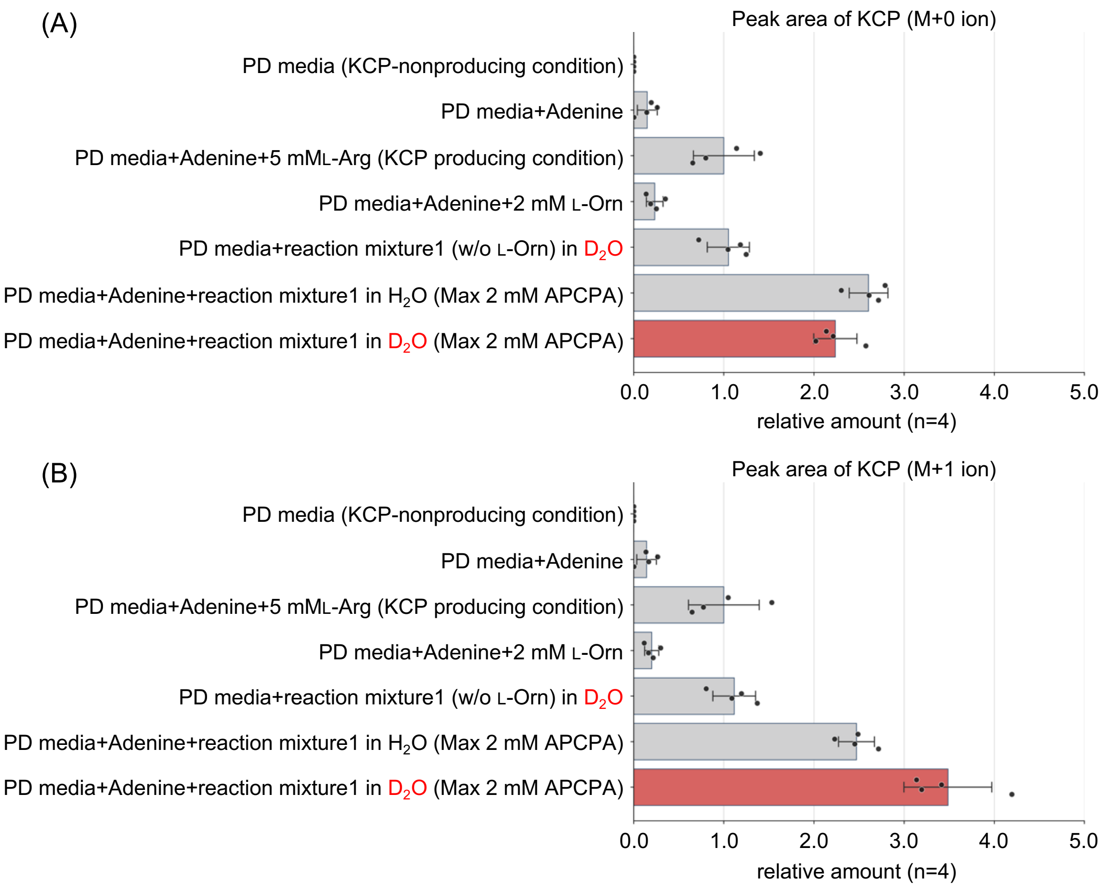


**Figure S12. Effect of reaction mixture feeding on KCP production.**

(A) Relative peak areas of protonated KCP M+0 ion under each culture condition. (B) Relative peak areas of the protonated KCP M+1 ion under each culture condition. Feeding of the $H_2O$-prepared ACPCA-containing reaction mixture increased the overall KCP signal, indicating stimulation of de novo KCP production from endogenous unlabeled precursors. Feeding of the $D_2O$-prepared ACPCA-containing reaction mixture increased the M+1 isotopologue signal more strongly, consistent with incorporation of deuterium-enriched ACPCA-derived material into KCP. Because feeding also stimulated total KCP production, concurrent de novo biosynthesis of unlabeled KCP likely diluted the observed M+1/M+0 enrichment. Data are shown as mean ± SD from independent biological replicates; dots indicate individual replicates. In both panels, the value for the KCP-producing condition (PD medium supplemented with adenine and 5 mM L-Arg) was set to 1.0.

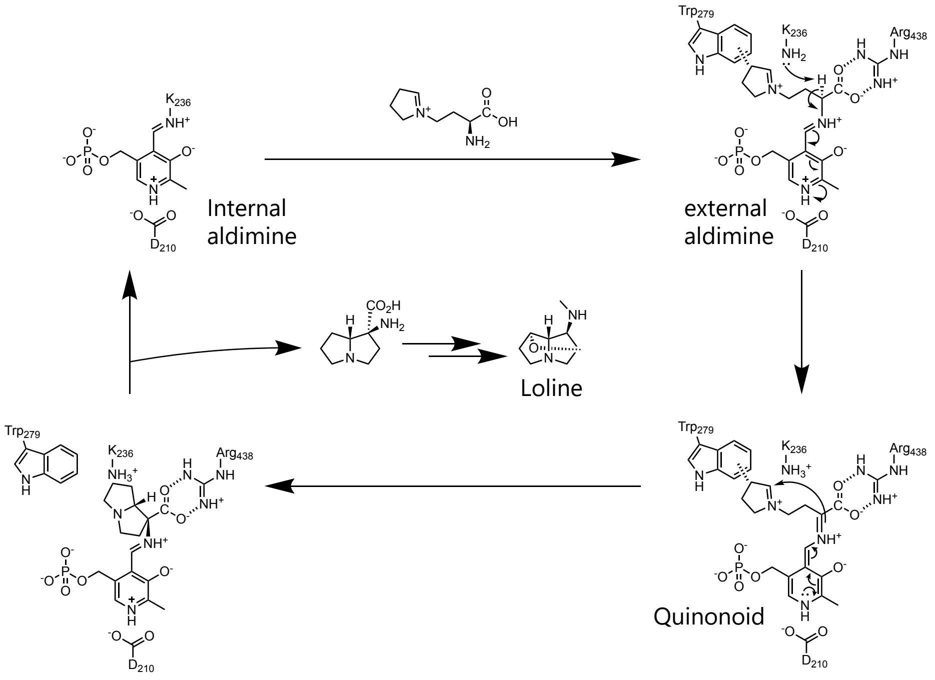


**Figure S13. Proposed catalytic mechanism of the PLP-dependent Mannich cyclase LolT.**

LolT employs pyridoxal 5′-phosphate (PLP) as a cofactor to catalyze an intramolecular Mannich-type C–C bond-forming reaction during loline alkaloid biosynthesis. In the resting state, PLP forms an internal aldimine with Lys236. Substrate binding and transaldimination generate the external aldimine, in which the substrate carboxylate is stabilized by Arg438 and the PLP cofactor is positioned through interactions including Asp210. Deprotonation at the α-position by Lys236 gives a PLP-stabilized quinonoid intermediate. The resulting carbanion/enamine then attacks the tethered iminium moiety to form the pyrrolizidine core. Trp279 is proposed to stabilize the iminium intermediate through a cation–π interaction and to help orient the substrate for stereoselective C–C bond formation. Subsequent protonation and product release regenerate the PLP–Lys236 internal aldimine. Adapted from the proposed mechanism of LolT reported by Gao et al.[5]

#### REFERENCES


(1) Jez, J. M.; Ferrer, J. L.; Bowman, M. E.; Dixon, R. A.; Noel, J. P. Dissection of Malonyl-Coenzyme A Decarboxylation from Polyketide Formation in the Reaction Mechanism of a Plant Polyketide Synthase. *Biochemistry* **2000**, *39* (5), 890–902.

(2) Maeno, Y.; Shiraishi, T.; Saito, N.; Maruyama, J.-I.; Shin-Ya, K.; Kuzuyama, T. Biosynthesis of Kaitocephalin: A Neuroprotective Natural Product Featuring a Peptide-like yet Nonpeptidic Scaffold. *Angew. Chem. Int. Ed Engl.* **2026**, No. e23010, e23010.

(3) Furihata, K.; Seto, H. J-Resolved HMBC, a New NMR Technique for Measuring Heteronuclear Long-Range Coupling Constants. *Tetrahedron Lett.* **1999**, *40* (34), 6271–6275.

(4) Hansen, P. E. Carbon—Hydrogen Spin—Spin Coupling Constants. *Prog. Nucl. Magn. Reson. Spectrosc.* **1981**, *14* (4), 175–295.

(5) Gao, J.; Liu, S.; Zhou, C.; Lara, D.; Zou, Y.; Hai, Y. A Pyridoxal 5′ -Phosphate-Dependent Mannich Cyclase. *Nat. Catal.* **2023**, *6* (6), 476–486.
